# Cell-nonautonomously tunable actomyosin flows orient distinct cell division axes

**DOI:** 10.1101/164186

**Authors:** Kenji Sugioka, Bruce Bowerman

**Affiliations:** Institute of Molecular Biology, 1229 University of Oregon, Eugene, Oregon 97403, USA.

**Author notes:** Corresponding author: Kenji Sugioka. Institute of Molecular Biology, University of Oregon, Eugene, OR 97403 USA.

**Keywords:** Mechanosensitivity, Embryogenesis, Dorsal-ventral axis, Cell division, Myosin, Cell polarity, Caenorhabditis elegans, Mouse

## Abstract

Cell division axes during animal development are arranged in diverse orientations, but the molecular mechanisms underlying this diversity remain unclear. By focusing on oriented divisions that are independent of known microtubule/dynein pathways, we show here that the non-muscle myosin II motor is an extrinsically tunable force generator that orients cell division axes through cortical actomyosin flows. We identified three extracellular cues that generate different actomyosin flows. A single contact site locally inhibited myosin activity in a mechanosensitive manner to generate local flow asymmetry, while size asymmetry of two contact sites and Wnt signaling both polarized myosin activity and actomyosin flow, with the latter overriding mechanosensitive effects. These intracellular actomyosin flow anisotropies specify distinct division axes to establish the geometries of not only *Caenorhabditis elegans* 4-, 6-, and 7-cell stage but also mouse 4-cell stage embryos. Tunable actomyosin flows together with microtubule/dynein pathways may specify diverse division axes across species.

## Highlights

Mechanosensitive actomyosin flow orients cell division independent of microtubules. Extrinsic Wnt signal abrogates mechanosensitive effects on cortical flow. Patterns of cell contact establish mouse and *C. elegans* embryonic geometries.

## Introduction

In animal development, cell division axes are arranged in diverse orientations relative to the niche or body axes during embryogenesis and stem cell division (Knoblich, 2010), and during brain (Egger et al., 2007), skin (Williams et al., 2014), kidney (Hao et al., 2010), and heart (Wu et al., 2010) organogenesis. Extrinsic controls of cell division orientation—whereby one cell instructs another to divide in a spatially organized manner—is a key mechanism of multicellular assembly (Siller and Doe, 2009; Knoblich, 2010; Williams and Fuchs, 2013; Rose and Gönczy, 2014). Consistent with this developmental significance, mutations in the genes required for oriented cell division are associated with various human diseases such as lissencephaly, microcephaly, leukemia, deafness, Huntington’s disease, and multiple cancers (Noatynska et al, 2012; Pease and Tirnauer, 2011). Although the molecular and physical mechanisms underlying oriented cell division have been studied extensively, these investigations have focused on a limited number of cell types. Extending such analyses to unexplored cells should expand our overall understanding of the molecular systems required to achieve division axis diversity *in vivo* and lead to the identification of cell division regulators that can be potential therapeutic targets.

In principle, upstream physical or chemical cues direct downstream force-generation machinery to orient the cell division axes. Thus far, the microtubule motor protein dynein is the only known force generator. Dynein works at two different cellular locations: the cell cortex and the cytoplasm. At the cell cortex, various cues—including cell polarity (Rose and Gönczy, 2014; di Pietro et al., 2016), tricellular junctions (Bosveld et al., 2016), or mechanical forces (Fink et al., 2011; Nestor-Bergmann et al., 2014)—localize an evolutionarily conserved protein complex composed of Gα, LGN, and NuMA. LGN/Gα/NuMA complex binds to dynein and generates microtubule pulling forces toward cell cortex via minus end-directed dynein movements, thereby orienting cell division axes to be perpendicular to the cortical site. In the cytoplasm, cell shape distortion acts as a cue that generates differences in astral microtubule length confined by the cell cortex. Longer astral microtubules then bind more cytoplasmic dynein to generate greater pulling force and hence orient division along the cell’s long axis (Minc et al., 2011), a mechanism also known as Hertwig’s rule (Wilson, 1925). Although recent reports suggest that F-actin and myosin motors also participate in cell division orientation, they modulate known pathways by controlling cell shape (Campinho et al., 2013) and NuMA localization (Seldin et al., 2013) or downstream microtubule pulling forces (Kwon et al., 2015). One study in *Caenorhabditis elegans* suggested that cell autonomous myosin motions tilt the division plane by approximately 20° in the clock-wise direction during establishment of the left-right body axis, but this model did not address the possible effects of cell shape or dynein pulling forces on spindle microtubules, or the upstream cues that influence myosin dynamics (Naganathan et al., 2014). Thus, microtubule motor dynein-dependent force generation machineries are the only known tunable mechanism of cell division orientation controlled by extracellular signals. However, fly and mouse without centrosome and astral microtubules still form a relatively normal body plan (Basto et al., 2006; Bazzi and Anderson, 2014), suggesting that pulling forces acting on astral microtubules may not be the sole tunable forces that drive oriented cell division.

Non-muscle myosin II is an actin-dependent motor protein that regulates cellular contractility and cortical flow—the concerted movement of a viscoelastic cell surface layer comprising F-actin, myosin, and cross-linking factors (Bray and White, 1988; Levayer and Lecuit, 2012). Cortical flow plays vital roles during development such as cell locomotion, growth cone migration, cell polarization, and cytokinesis (Bray and White, 1988). When a molecular clutch physically connects cortical actomyosin to the plasma membrane, cortical flow generates forces that drive cell migration or morphogenesis (Case and Waterman, 2015; Roh-Johnson et al., 2012). The mechanistic basis for cortical flow orientation is the anisotropy of cellular contractility, which generates flow from regions of low to high contractility within the cell (Mayer et al., 2010). The velocity of cortical flow is further regulated by actin dynamics and non-muscle myosin II motor activity (Mayer et al., 2010). In the case of cytokinesis, cortical flow is oriented toward the equatorial region of the cell and contributes to contractile ring assembly (Uehara et al., 2010; Yumura et al., 2008; Zhou and Wang, 2008; Reymann et al., 2016). Inhibiting myosin activity or actin dynamics limits flow velocity, resulting in cytokinesis failure (Murthy and Wadsworth, 2005). Most studies to date have focused on single cells, while the regulation of cortical flow in multicellular system and its role during cell division remain poorly understood.

In the present study, we identified an extrinsically tunable myosin-dependent force generation mechanism that controls cell division orientation in both *C. elegans* and mouse embryos. In multicellular contexts, a single cell may receive multiple physical and chemical cues from different neighboring cells, complicating the identification of critical cues. To overcome this predicament, we used isolated blastomeres and adhesive polystyrene beads to reconstitute multicellular environments. We identified three extracellular cues that control cortical actomyosin flow during oriented cell divisions: physical contact, asymmetrical contact size, and Wnt signaling. Each of these cues modified myosin activity to generate distinct cortical flows that differently oriented cell divisions to establish multicellular architecture in *C. elegans* 4, 6, and 7-cell stage embryos. Moreover, we show that a similar mechanosensitive mechanism oriented cell division in mouse embryos, and document a conserved physical basis underlying the establishment of 4-cell stage architectures in both nematodes and mice, and likely also in humans. Our discovery of a novel molecular mechanism that regulates the orientation of cell division axes significantly advances our understanding of division axis diversity and the assembly of multicellular architectures during animal development.

## Results

### Oriented AB cell division during C. elegans dorsal-ventral (D-V) axis establishment is a microtubule-independent process

To identify oriented cell division mechanisms that are independent of known microtubule/dynein pathways, we focused on AB cell division in 2-cell stage *C. elegans* embryos. The anterior AB cell divides before the posterior P_1_ cell, and their daughters always adopt a diamond shape at the 4-cell stage, with the posterior daughter of AB (ABp) and anterior daughter of P_1_ (EMS) aligned perpendicularly to the anterior-posterior (A-P) axis to define the D-V body axis (Fig. 1A) (Priess and Thomson, 1987; Sulston and Horvitz, 1977). This diamond shape is critical for later development, as signals from P_2_ activate Notch signaling in ABp and Wnt signaling in EMS to specify different cell fates (Fig. 1A) (Priess, 2005; Sawa and Korswagen, 2013). During cell division, the AB and P_1_ mitotic spindles are oriented parallel and perpendicular to the plane of AB-P_1_ cell contact, respectively (Fig. 1B). P_1_ spindle orientation is regulated by an upstream cue from the midbody remnant of zygotic division and a downstream LGN/dynein-dependent force generation mechanism (Singh and Pohl, 2014; Srinivasan et al., 2003). However, the AB division axis oriented normally after LGN knock-down, suggesting that oriented AB division is independent of cortical dynein-dependent microtubule pulling forces (Fig. 1B). Furthermore, the long axis of the AB cell before cell division did not correlate with its division axis, indicating that the oriented AB division is regulated independently of Hertwig’s rule (Fig. S1A-B). To assess the potential roles of other microtubule-dependent mechanisms in AB cell division orientation, we treated embryos with the microtubule-depolymerizing drug nocodazole (12.5 ng/ml) and found that the division axes of AB were unaffected while those of P_1_ were randomized (Fig. S1C). Furthermore, even when spindle formation was abolished by treatment with 20 μg/ml nocodazole, non-muscle myosin II still formed a cleavage furrow in AB that was oriented perpendicularly to the plane of cell contact (Fig. 1C). These results indicate that a previously unknown microtubule-independent mechanism establishes the AB division axis.

**Figure 1.**
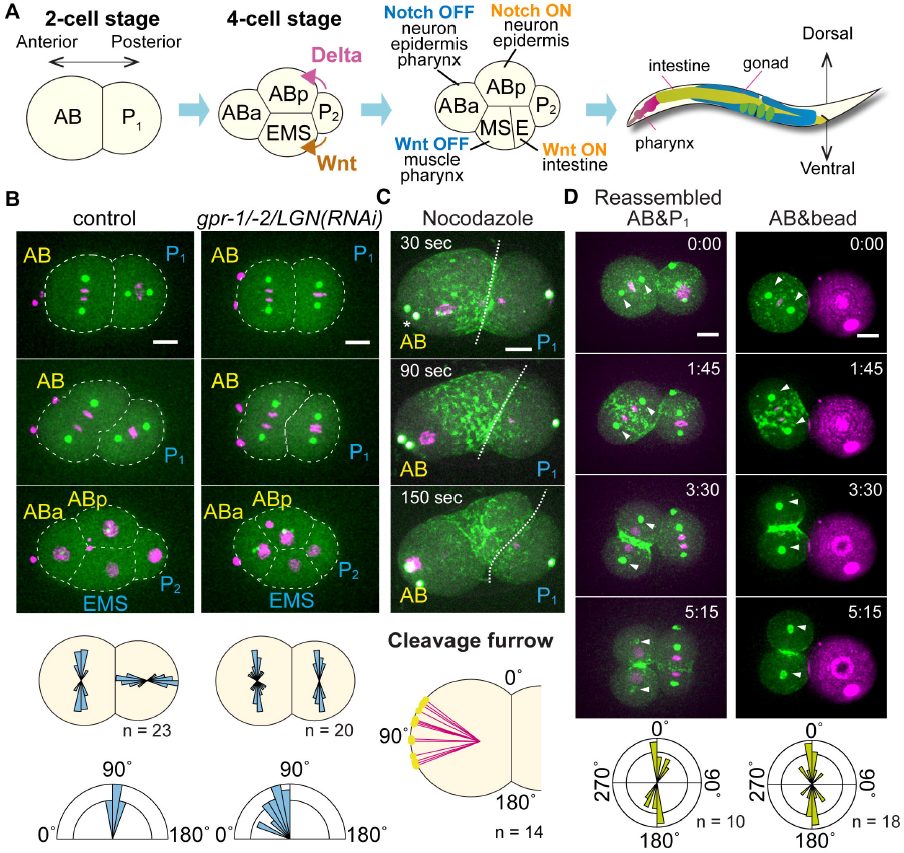
A microtubule-independent mechanosensitive mechanism orients AB blastomere division during D-V body axis establishment. (A) The 4-cell stage architecture establishes the *C. elegans* D-V body axis (See text). (B) Orientation of AB and P_1_ cell divisions. Centrosomes (green), histone H2B (magenta), and cell outlines (white dotted line) are shown along with the distribution of division axes (middle panels) and D-V axis orientation (bottom panels). (C) Myosin accumulation in the nocodazole-treated cells. Non-muscle myosin II (green), centrosomes (white), and histones (magenta) are shown along with cleavage furrow position (bottom). (D) Orientation of AB cell division after attachment to P_1_ cells or beads. Myosin (green), centrosomes (green), histones (magenta), and beads (magenta) are shown along with the distribution of division axes (bottom). Scale bars = 10 μm.

### Myosin activity regulates AB cell division orientation

We investigated the mechanisms underlying oriented AB cell division. We found that knockdown of the Cullin E3 ubiquitin ligase component CUL-3 resulted in a randomization of the AB division axis, with 27% of the 4-cell stage embryos showing a linear cell arrangement and thus a severely disrupted D-V axis (Fig. 2A). During cell division in *cul-3(RNAi)* embryos, non-muscle myosin II foci were abnormally distributed throughout the cell cortex, although a functional contractile ring was subsequently formed (Fig. 2B, D), suggesting that CUL-3 acts via an actomyosin-dependent pathway to orient AB cell division. To evaluate this possibility, we manipulated non-muscle myosin II activity by pharmacological treatment or RNAi. The Rho GTPase-activating protein RGA-3 targets RhoA and its knock-down activates myosin II (Fig. 2C) (Schonegg et al., 2007). We found that *rga-3(RNAi)* resulted in broader distribution of myosin II foci and abnormal orientation of AB division axes, similar to what we observed after *cul-3(RNAi)* (Fig. 2B). ML-7 is a myosin light chain kinase inhibitor that inactivates myosin II (Fig. 2C) (Saitoh et al., 1987); ML-7 treatment inhibited cell contraction and randomized the orientation of AB division axes (Fig. 2F). Taken together, these results indicate that myosin II activity regulates orientation of the AB division axis.

**Figure 2.**
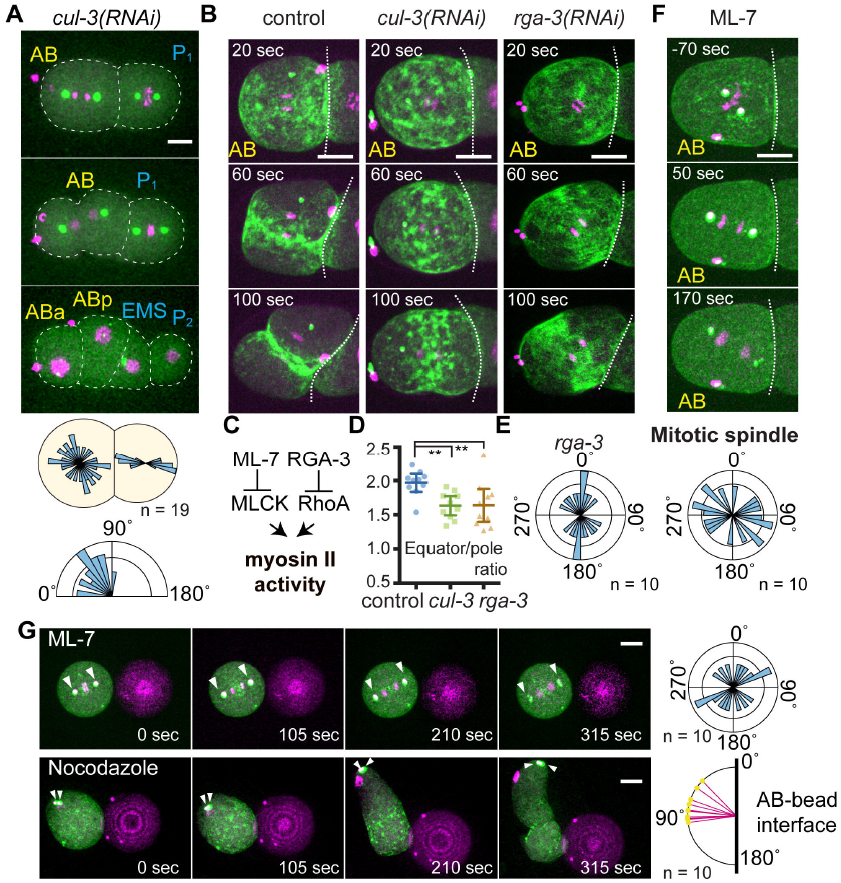
Myosin activity regulates mechanosensitive AB division axis orientation. (A) Abnormal AB division axis in *cul-3*(RNAi). Centrosomes (green), histone H2B (magenta), and cell outlines (white dotted lines) are shown along with the distribution of division axes (middle panel) and D-V axis orientations (bottom panel). (B) Myosin localization during AB cell division. Myosin (green) and histones (magenta) are shown. (C) Diagram of myosin II regulation. (D) Ratio of equatorial to polar cortical myosin intensity. P values were calculated by one-way ANOVA with Holm-Sidak’s multiple comparisons test. **P < 0.01. (E) AB division axis orientations in *rga-3*(RNAi). (F) AB division axis orientation after ML-7 treatment. Myosin (green), centrosomes (white), and histones (magenta) are shown along with the distribution of mitotic spindle orientations (bottom). (G) Orientation of AB division relative to the bead after drug treatment. Myosin (green), centrosomes (white), histones (magenta), and beads (magenta) are shown; arrowheads indicate centrosome positions. Distributions of mitotic spindle orientations or cleavage furrows are shown in the right panels. Scale bars = 10 μm.

### Physical contact is an upstream cue for AB cell division orientation

We next explored the origin of upstream cues that orient AB cell division. To test whether cell contact serves as a cue, we isolated early 2-cell stage AB and P_1_ cells and recombined them in culture medium to create contact sites at random locations (Fig. 1D). The AB division axis was initially randomly oriented, but then rotated to become aligned with the AB-P_1_ boundary by the end of telophase (hereafter referred to as parallel division) (Fig. 1D and Movie S1), suggesting that cell contact is an upstream cue that orients AB cell division. Extrinsic cues transmitted through cell contact can be either mechanical or chemical in nature. To evaluate the former possibility, we used carboxylate-modified polystyrene beads that nonspecifically bind to the amine groups of extracellular proteins. Attachment of a bead to isolated AB cells resulted in parallel division (Fig. 1D and Movie S1), demonstrating that physical contact is sufficient to orient AB cell division. To examine whether physical contact distorts cell shape, we measured the perpendicular and parallel cell diameters relative to the beads and found that their ratio was approximately 1.0 (mean ± SD: 0.991 ± 0.021, n = 20), indicating that cell-bead contact did not cause changes in cell shape and that bead-induced oriented AB division was independent of Hertwig’s rule. Thus, mechanical signals from cell contact are an upstream cue that directly orients AB cell division.

We also investigated the requirement for myosin activity in the mechanosensitive regulation of division axis orientation. ML-7 treatment in the cell-bead experiment caused randomization of the AB division axes relative to the beads (Fig. 2G, upper panel). In contrast, neither nocodazole treatment nor LGN knockdown affected cleavage furrow orientation (Fig. 2G, bottom panel and Fig. S2B). Although cytokinesis was defective following ML-7 treatment, this was not the direct cause of abnormal division axis orientation, as cytokinesis-defective cells from ZEN-4/MKLP-1 mutants oriented normally in parallel to the bead (Fig. S2B). Taken together, these data indicate that physical contact orients AB cell division through an actomyosin-dependent mechanosensitive process.

### Physical contact induces intracellular myosin flow anisotropy

To understand how physical contact controls actomyosin activity, we analyzed the dynamics of cortical myosin II foci. We defined the x axis as passing through the spindle poles and the y axis as perpendicular to the x axis and within the plane of the cell and bead centers (Fig. 3A). We also divided the AB cell surface into four quadrants–– U1, U2, B1, and B2––with B2 attached to the bead (Fig. 3A). Cell movements were computationally removed to precisely analyze cortical myosin II dynamics. In the absence of a bead, myosin foci flowed toward the cell equator along the x axis and exhibited no directional y-axis movements (Figs. 3A, B and S4 and Movie S2). However, upon bead attachment, x-axis myosin movements were significantly reduced in B2 (Figs. 3A, B and 4A and Movie S2, S3), and y-axis movements were observed that were always in the counter-clockwise direction around the division axis when viewed from the nearest pole, with U1 and B1 moving in the direction opposite to U2 and B2 (Figs. 3A and S4 and Movie S2). Intact embryos exhibited similar myosin movements (Fig. 3A–C), suggesting that contact with either beads or cells induces anisotropic myosin II flow. In *cul-3(RNAi)* embryos, where division axes became abnormal, x-axis myosin flow asymmetry was lost, whereas y-axis movements were similar to the control (Fig. 3A-C), suggesting that the localized x-axis myosin flow asymmetry is important for oriented cell division. Moreover, although y-axis myosin movements were always counter-clockwise regardless of bead position, the x-axis myosin movements were specifically reduced at the contact site. Due to this spatial response, the equatorial myosin flows distal and proximal to the contact site were symmetric and asymmetric, respectively, creating a local myosin flow asymmetry within the cell (Fig. 7G, black arrows in the left panel and Movie S3).

**Figure 3.**
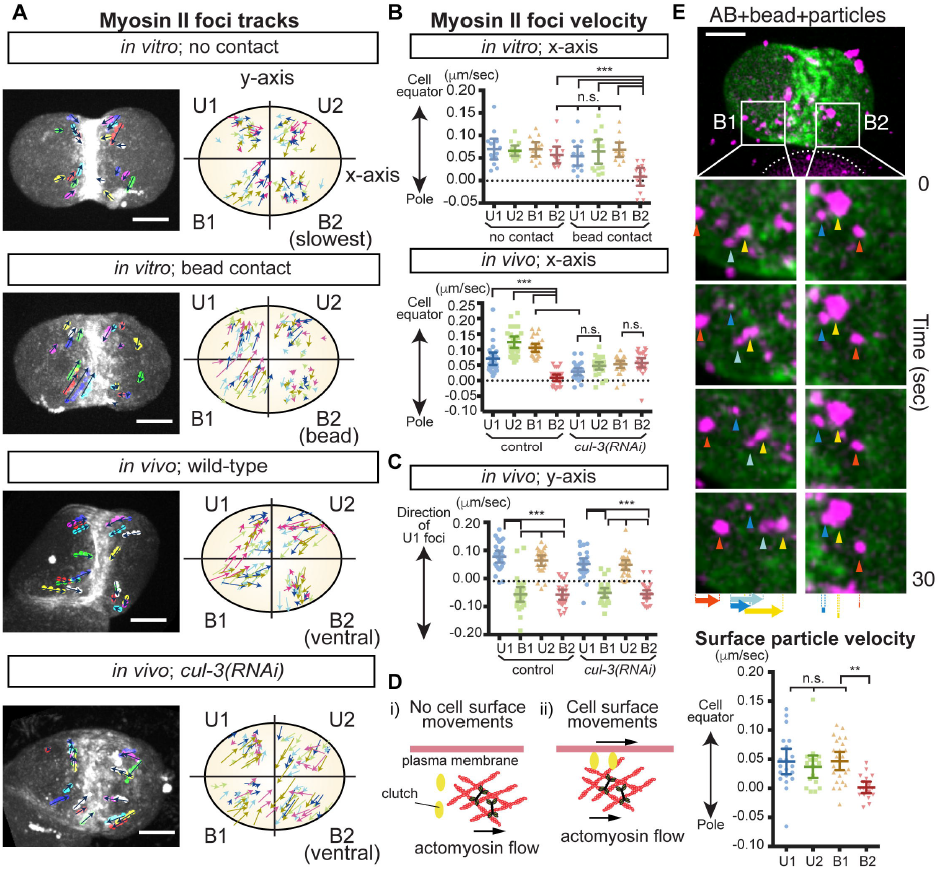
Mechanosensitive induction of anisotropic actomyosin flow triggers cell surface movements. (A) Movements of non-muscle myosin II/NMY-2 foci during AB cell division. Isolated AB cells with or without bead attachment and control or *cul-3*(RNAi) embryos. Myosin foci tracks for 50 s are shown in left panel. In the right panel, color of arrows indicate the tracks from different samples. The cell quadrant was defined as indicated. In the cells with no contact, regions exhibited slowest myosin velocities are B2 for convenience of comparison. (B, C) Velocities of NMY-2 foci in x and y axes of the cell defined in A. (D) Clutch engagement transmits flow forces to the cell surface (see text). (E) Cell surface movements during oriented AB division. Movements of 0.35-μm particles attached to the membrane (arrowheads) and their velocities are shown along with myosin (green) and beads (white dotted line). P values were calculated by one-way ANOVA with Holm-Sidak’s multiple comparisons tests. Scale bars = 10 μm.

### Physical contact limits RLC phosphorylation to generate anisotropic actomyosin flow

We next investigated how bead attachment or cell contact limits equatorial (x-axis) myosin flow. First, the velocity of equatorial myosin flow in B2 was inversely proportional to bead diameter (Fig. 4A, B). In addition, the duration of myosin foci on the cell cortex (i.e., foci lifetime) was reduced at the contact site (Fig. 4D), implying that physical contact controls myosin II activity.

**Figure 4.**
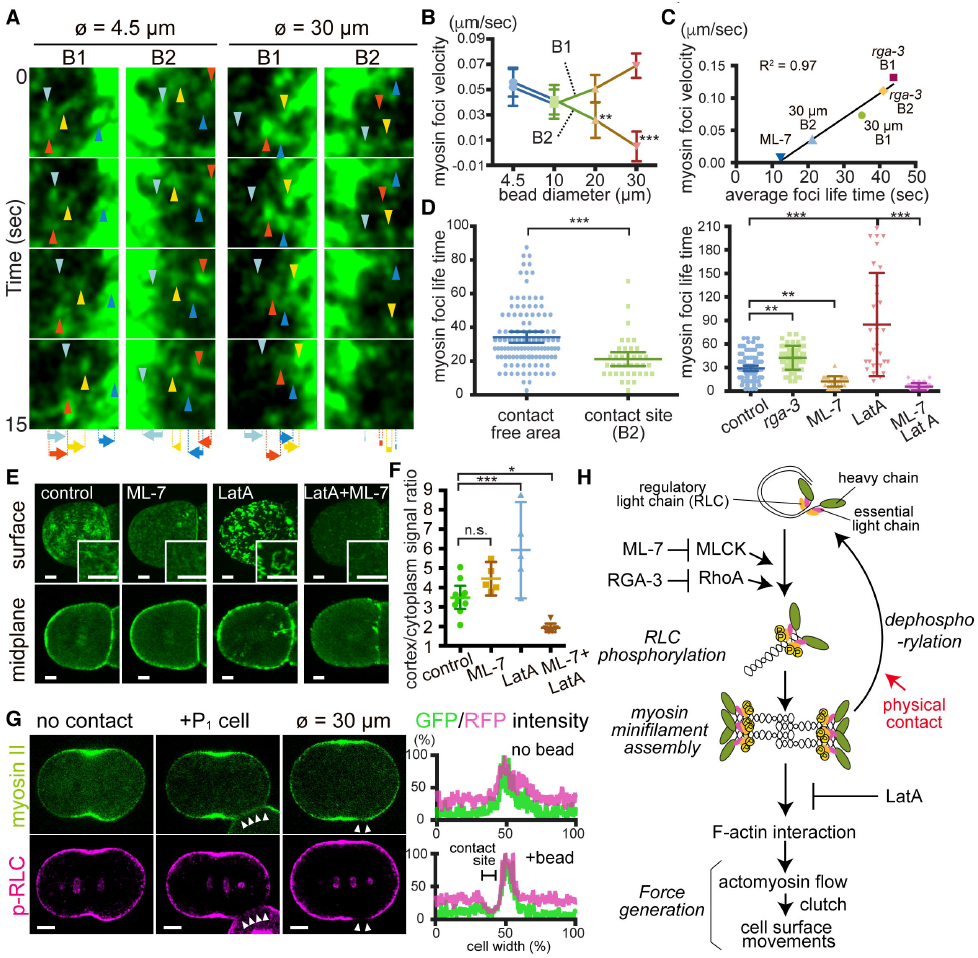
Mechanosensitive inhibition of myosin RLC phosphorylation generates anisotropic actomyosin flow. (A) Myosin foci movements during oriented AB division in response to beads of different sizes. Arrowheads and arrows indicate myosin foci and their total displacement, respectively. (B) Relationship between myosin foci velocity and attached bead diameter. (C) Relationship between myosin foci velocity and myosin foci lifetime. (D) Myosin foci lifetime in response to beads or different myosin activities. (E) p-RLC-dependent myosin foci formation. (F) Cortex-to-cytoplasm ratio of myosin intensity. (G) Contact-dependent changes in p-RLC localization. Arrowheads indicate the cell or bead contact site. (H) Pathway diagram of contact-dependent myosin flow control (see text). Scale bars = 10 μm. P values were calculated by one-way ANOVA with Holm-Sidak’s multiple comparisons test. *P < 0.05, **P < 0.01, ***P < 0.0001; n.s., not significant (P > 0.05).

We thus sought to position the role of physical contact within the cascade of biochemically characterized myosin regulatory pathways (Fig. 4H). For non-muscle myosin, each myosin II heavy chain dimer binds two essential light chains and two regulatory light chains (RLC) to form a hexamer. RLC phosphorylation at the evolutionarily conserved Thr 18 and Ser 19 residues controls myosin activity: the ATPase activity of myosin is increased by the RLC phosphorylation (Adelstein and Conti, 1975) and RLC phosphorylation triggers a conformational change from a closed to an open form that allows minifilament assembly and promotes downstream contraction (Fig. 4H) (Craig et al., 1983; Scholey et al., 1980; Vicente-Manzanares et al., 2009). We first assessed whether the myosin foci we observed *in vivo* represent myosin minifilaments. When we blocked F-actin assembly using the actin-depolymerizing drug latrunculin A (LatA), myosin foci were stabilized with some exhibiting a filamentous morphology (Fig. 4D, E), suggesting that myosin foci formation does not require F-actin interaction. On the other hand, inhibition of RLC phosphorylation by ML-7 treatment resulted in the loss of myosin foci and reduced foci life time (Fig. 4D, E). As ML-7 treatment did not affect cortical localization of myosin, reduced foci lifetime suggests they disassemble rather than detach from the cortex (Fig. 4E, F). Furthermore, activation of RLC phosphorylation by *rga-3(RNAi)* increased foci life time (Fig. 4D). These results suggest that myosin foci are myosin minifilaments with their life time regulated by RLC phosphorylation. Notably, the level of RLC phosphorylation correlates with myosin foci velocity (Fig. 4C). Therefore, mechanosensitive reduction of equatorial myosin flow velocity at the contact site may be caused through the inhibition of RLC phosphorylation.

To determine whether RLC phosphorylation is indeed reduced by cell or bead contact, we performed immunolabeling using anti p-RLC (Ser19) antibody that recognizes *C. elegans* MLC-4/RLC (Zonies et al., 2010). In the absence of contact, p-RLC co-localized with myosin II-GFP (Fig. 4G). On the other hand, after P_1_ cell or bead attachment, p-RLC signal intensity at the contact site was markedly reduced to 57 ± 6% of the intensity at the pole region (P = 0.0012), in contrast to myosin-GFP (136 ± 15% intensity relative to the pole region) (Fig. 4G). Thus physical contact locally inhibits myosin RLC phosphorylation, thereby limiting myosin activity and flow during the generation of local actomyosin flow asymmetry (Fig. 4H).

### Actomyosin flow generates forces to trigger cell surface movements

For the control of cell division orientation, actomyosin flow needs to generate forces that act on the cell surface. To determine whether actomyosin flow generates forces that trigger cell surface movement, we coated isolated AB cells attached to a 30 μm bead with carboxylate-modified particles (0.35 μm in diameter) and found that the surface particles exhibited flows that were similar to those of myosin II foci and also were limited near the contact site (Fig. 3E and Movie S4). These results suggest that mechanosensitive actomyosin flow generates forces to move the cell surface through a molecular clutch that links the actomyosin cortex to the plasma membrane (Fig. 3D) (Case and Waterman, 2015). Given that equatorial cell surface movement is asymmetric in B1 and B2 but not in U1 and U2, we propose that cell surface proximal to the contact site moves toward B2, generating directional torque and orienting the dividing cell in parallel to the contact site (Fig. 7G, orange arrows in the left panel).

### Mechanosensitive myosin pathway orients cell division in C. elegans and mouse embryos to establish the 4-cell architecture

We investigated whether the mechanosensitive myosin-dependent mechanism we described also operates in mammals using mouse embryos. As in *C. elegans*, the 2-cell stage mouse embryo undergoes asynchronous divisions; the early dividing cell is called AB and the other CD (Kelly et al., 1978). Both AB and CD cells in intact 4-cell stage embryos underwent division in parallel to the contact site; 88% of embryos had a tetrahedral shape and 12 % were diamond shaped (Fig. 5A, C and Movie S5). Previous studies have shown that the mitotic spindle was randomly oriented at metaphase (Louvet-Vallee et al., 2005) and removal of the zona-pellucida—a glycoprotein layer surrounding cells—resulted in abnormal cell division orientation (Graham and Deussen, 1978; Suzuki et al., 1995), implying that steric confinement by the zona pellucida, rather than active force generation, orients these cell divisions. However, we found that in embryos lacking the zona pellucida (i.e., zona-free embryos), AB division always oriented in parallel to the contact site (Fig. 5A and Movie S5). We therefore assessed whether AB/CD cells respond to physical contact. When attached to beads, isolated AB or CD blastomeres underwent parallel division, suggesting that physical contact acts as a cue for oriented division (Fig.5A and Movie S5). Moreover, ML-7 treatment inhibited parallel division (Fig. 5A) and cell or bead contact locally reduced p-RLC localization (Fig. 5B and Fig. S5). Thus, as in *C. elegans*, mouse 2-cell stage embryos undergo mechanosensitive myosin-dependent oriented cell division.

**Figure 5.**
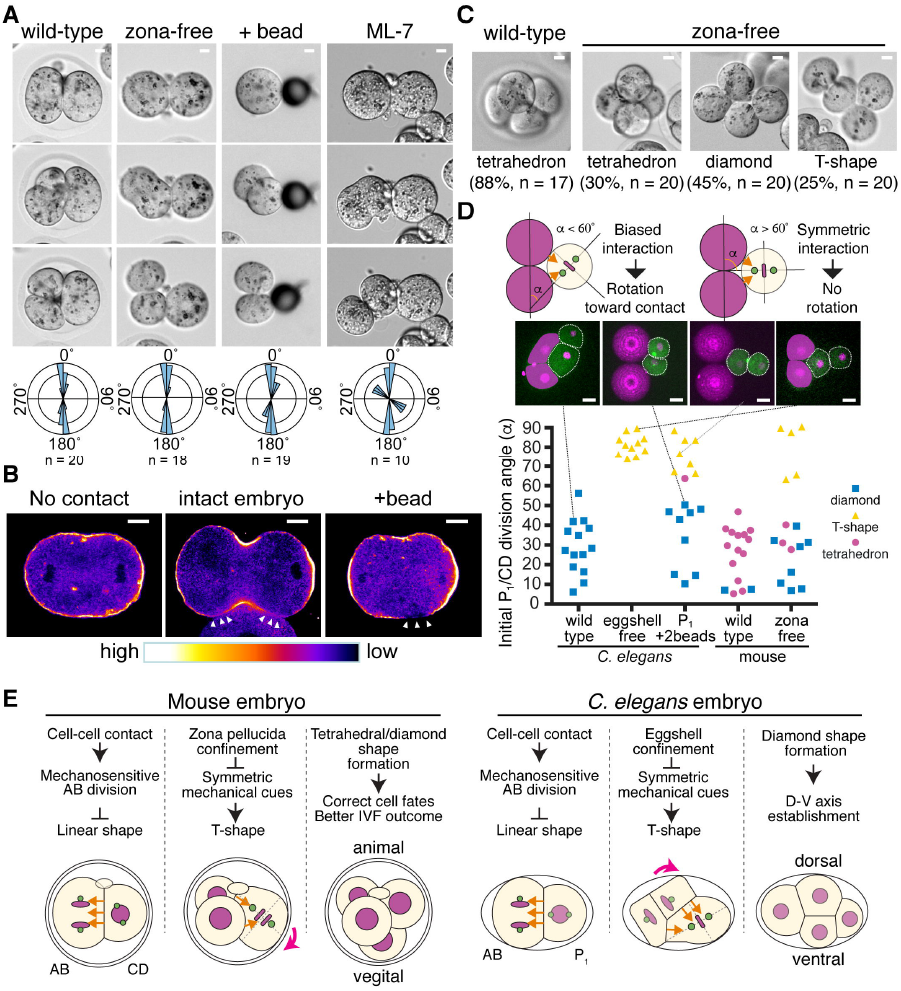
Mechanosensitive myosin pathway orients cell division in *C. elegans* and mouse embryos to establish 4-cell stage architecture. (A) Oriented AB cell division in mouse 2-cell stage embryos. The distributions of division axes is shown at the bottom. (B) Contact-induced changes in p-RLC localization. p-RLC localizations are shown as heat map. Arrowheads indicate contact sites. (C) 4-cell architecture after CD cell division. (D) Effects of initial P_1_ (metaphase) and CD (onset of cell elongation) division axis orientation and extracellular material on multicellular architecture formation. α, Angle of P_1_/CD division relative to the contacting cell. (E) Common mechanism underlying mouse and *C. elegans* 4-cell architecture formation (see text). Scale bars = 10 μm.

We next investigated how the mechanosensitive myosin-dependent pathway contributes to the establishment of *C. elegans* and mouse 4-cell embryo architectures that are critical for later development. As described above, previous study has indicated that zona-free mouse embryos exhibit abnormal division axes and 4-cell stage architecture. We confirmed these defects in zona-free embryos but in a more specific way; 25% of CD divisions were perpendicular to the contact site, resulting in T-shaped 4-cell embryos (Fig. 5C and Movie S6). These results do not indicate that CD cells are less capable of parallel division; indeed isolated CD cells always responded to bead contact with parallel divisions (Fig. 5A). Rather, the difference between the AB and CD divisions within embryos is their number of contact sites, with AB having one contact with CD and CD having two contacts, one with each AB daughter.

We analyzed the significance of two contacts and extracellular material on cell division orientation in mouse and *C. elegans*, by using wild-type embryos, embryos without extracellular materials (eggshell-or zona-free), and isolated *C. elegans* P_1_ cells attached to two beads (Fig. 5D). As in the case of CD cells, P_1_ cells attached to a single bead always underwent parallel division (Fig. S1D). However, when CD or P_1_ have two contact sites, we found that the angle of initial cell division (at metaphase in *C. elegans* and at the onset of cell elongation in mouse), relative to a line passing through the centroids of the two contacting beads or cells (defined as α), was the major determinant of the division pattern (Fig. 5D). When α was < 60°, P_1_ and CD cells rotated toward contact sites to undergo parallel division (Fig. 5D). However, this rotation did not occur when α was > 60°, possibly due to symmetric contacts with the B1 and B2 regions, and most cells formed T-shaped patterns (Fig. 5D). Additionally, α was < 60° in both *C. elegans* and mouse in the presence of extracellular material, suggesting that steric confinement affects the initial P_1_/CD division angle (Fig. 5D). Indeed, the eggshell/zona pellucida ensures the rotation of dividing AB cells in *C. elegans* or CD cells in mouse toward the cell contact (Movie S6). These results suggest that extracellular material prevents the selection of an alternative division axis created by the additional cell contact, thereby ensuring parallel P_1_/CD cell division in intact embryos. Finally, the difference in diamond/tetrahedral shape distribution between *C. elegans* and mouse 4-cell stage are likely due to the ellipsoidal and spherical extracellular material shapes, respectively (Yamamoto and Kimura, 2017). Thus, the 4-cell architecture in mouse and *C. elegans* is established via a common sequence of events: physical contact-dependent parallel division orients AB cell division, and extracellular material-dependent steric hindrance prevents P_1_/CD cell from having symmetric contacts with AB daughters that would otherwise lead to a T-shaped architecture (Fig. 6C).

**Figure 6.**
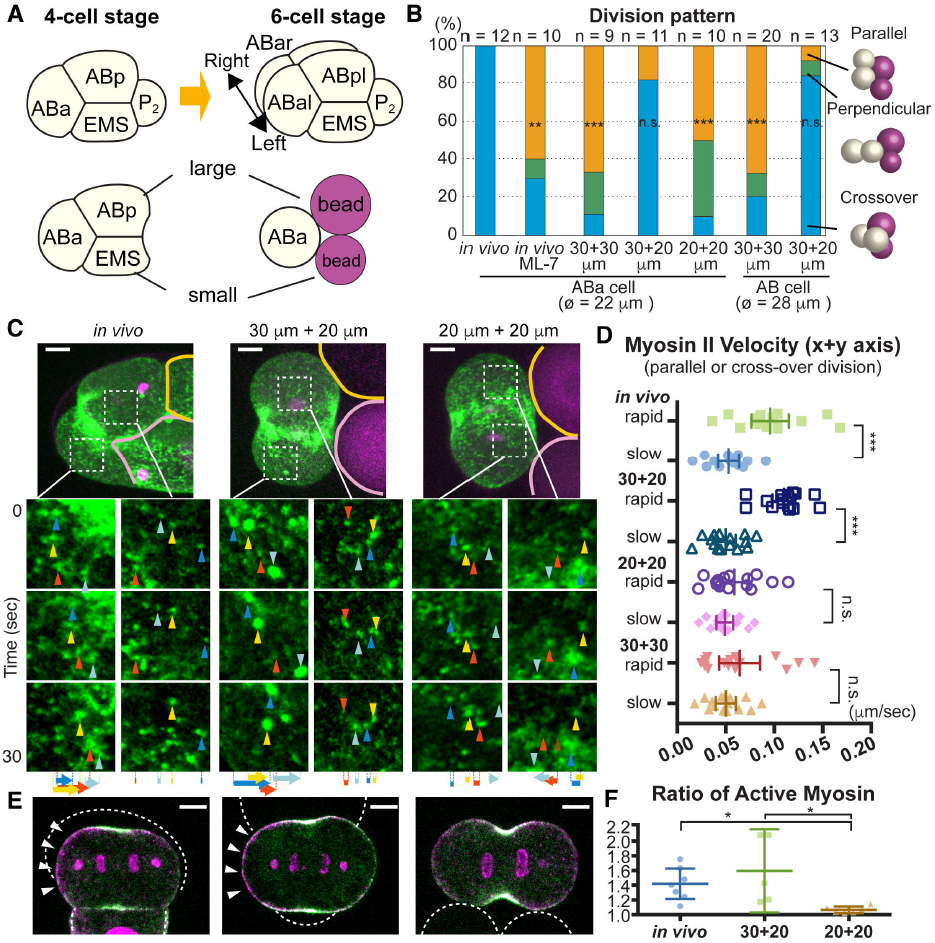
Asymmetry in cell contact size polarizes acomyosin flow to orient left-right division. (A) ABa cell division at the 4-cell stage and recapitulation of contact asymmetry using beads. (B) Effects of contact size asymmetry on ABa cell division axis. (C, D) Myosin foci movements during ABa cell division. Arrowheads and arrows in C indicate myosin foci and their total (A) displacement, respectively. Velocities are shown in D. (E) Polarized p-RLC localization in response to asymmetric-sized contacts. Myosin (green), p-RLC (magenta), cell or bead outlines (white dotted lines), and polarized p-RLC (arrowheads) are shown. (F) Ratio of active myosin intensities between opposite halves of cells. Active myosin intensity was determined as the ratio of p-RLC to myosin intensity. Scale bars = 5 μm. P values in B were determined with the Fisher’s exact test and those in D and F were calculated by one-way ANOVA with Holm-Sidak’s multiple comparisons test. *P < 0.05, **P < 0.01, ***P < 0.0001; n.s., not significant (P > 0.05).

### Second cue: asymmetric-sized contacts polarize actomyosin flow to orient left-right division axis

We also investigated myosin-dependent division axis control in later-stage *C. elegans* embryos. After formation of the 4-cell stage diamond shape, ABa cells divide along the left-right body axis (Fig. 6A) (Sulston et al., 1983). ML-7 treatment disrupted ABa cell orientation, suggesting that this process requires actomyosin activity (Fig. 6B). Before cell division, ABa is adjacent to larger ABp and smaller EMS cells, creating asymmetric-sized contacts. To recapitulate this situation, isolated ABa cells were attached to asymmetric (30-and 20-μm) beads. Upon attachment, 82% of ABa cells underwent a “crossover”-type division analogous to the left-right oriented division in wild-type embryos (Fig. 6B and Movie S7). In contrast, when ABa cells were attached to symmetric-sized (two 30--μm or two 20--μm) beads, the frequency of the crossover-type division was decreased (Fig. 6B). AB cells—the mother of ABa with larger size—showed similar behaviors (Fig. 6B). These results suggest that asymmetric cell contacts serve as a cue that induces crossover-type cell division, which in an intact embryo results in left-right oriented ABa cell division.

Analysis of myosin foci velocities revealed that both *in vivo* or upon attachment of asymmetric-sized beads, the two halves of the ABa cell showed asymmetric myosin velocities (Fig. 6C, D and Movie S7). Conversely, attachment to symmetric-sized beads yielded symmetric velocities (Fig. 6C, D and Movie S7). Asymmetric-sized contacts also induced a polarized localization of p-RLC, suggesting an asymmetric activation of myosin in these dividing cells (Fig. 6E, F). These results indicate that asymmetry in cell contact size is a cue that polarizes myosin activity and flow to specify left-right oriented ABa cell division.

### Third cue: extrinsic Wnt signal polarizes actomyosin flow to abrogate mechanosensitive effects

Following ABa cell division at the 6-cell stage, the EMS cell receives a Wnt signal from the posteriorly located P_2_ cell and undergoes asymmetric cell division to produce anterior mesoderm and posterior endoderm precursor cells (Fig. 7A) (Rocheleau et al., 1997; Thorpe et al., 1997). Despite EMS cell contacts with ABal, ABar, ABpl, ABpr, and P_2_ cells, EMS division is oriented toward P_2_ in a Wnt-dependent manner (Schlesinger etl al., 1999; Goldstein et al., 2006). We therefore investigated how EMS avoids mechanical influences from cells other than P_2_. When attached to a single bead, EMS cells underwent parallel division (Fig. S1D). In addition, when attached to a bead on one side and to ABxx cells (i.e., a pair of ABa or ABp daughters) on the other, EMS cells still underwent parallel division relative to the bead (Fig. 7B). However, when attached to a bead on one side and P_2_ on the other, EMS division axes were randomized relative to the bead (Fig. 7B and Movie S8). Furthermore, EMS cells attached to a bead on one side and P_2_ isolated from Wnt mutants on the other underwent parallel division (Fig. 7B and Movie S8). These results suggest that contact with P_2_ cancels the mechanical influence on cell division axes in a Wnt-dependent manner.

**Figure 7.**
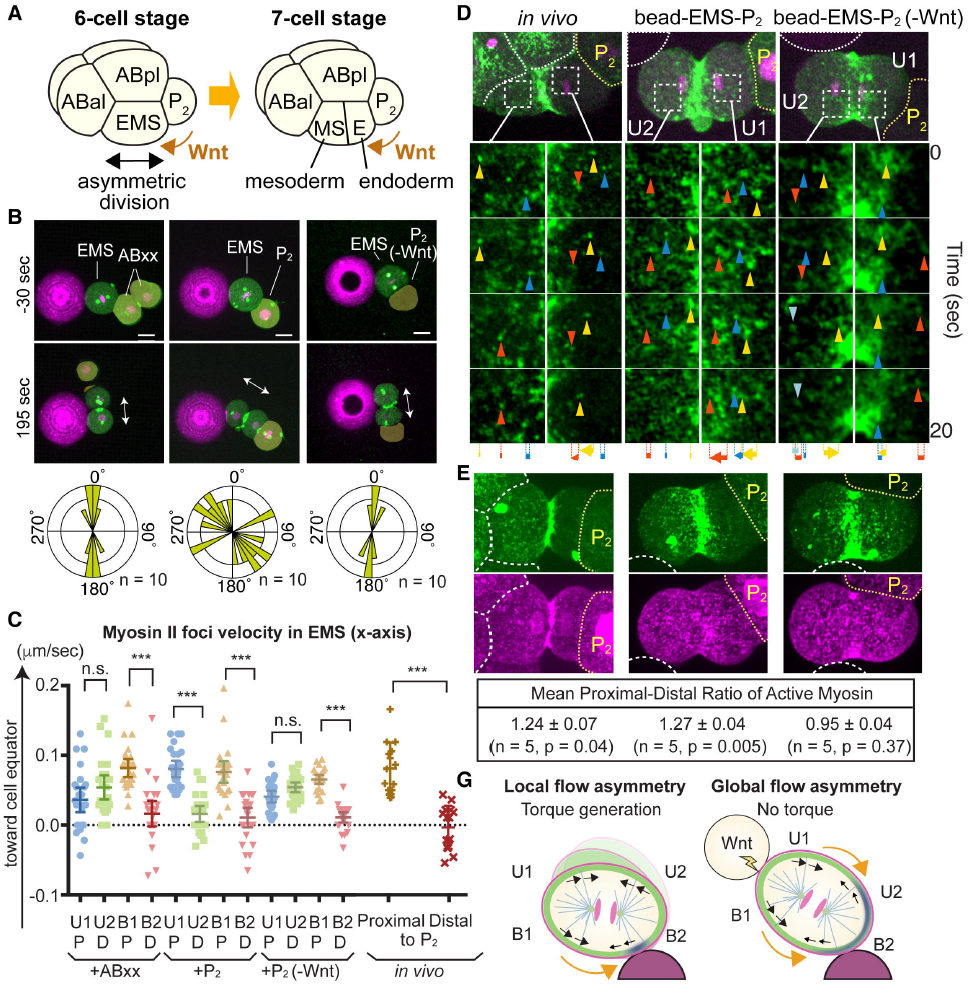
Extracellular Wnt signal polarizes actomyosin flow that abolishes mechanosensitive effects. (A) Asymmetric EMS division at the 6-cell stage is oriented toward P_2_ cells in the presence of multiple contacts. (B) Ignorance of beads contact induced by a Wnt signal from P_2_ cell. Myosin (green), centrosomes (green), histones (magenta), beads (magenta), and EMS division axis (arrows) are shown. P_2_(-Wnt) indicates the P_2_ cell isolated from Wnt mutants. (C, D) Myosin foci velocities (C) and movements (D) during EMS division. Arrowheads and arrows indicate myosin foci and their total displacement, respectively. (E) Wnt-dependent polarization of active myosin localization. Myosin (green), p-RLC (magenta), cell or bead boundaries (white dotted lines), and P_2_ cells (yellow dotted lines) are shown. Proximal-distal ratio of active myosin calculated similarly to Figure 6F is shown at the bottom. (G) Model of local actomyosin flow asymmetry oriented division and its abrogation by global flow asymmetry. Myosin (black arrows) and cell surface (orange arrows) movements are shown. P values were determined by one-way ANOVA with Holm-Sidak’s multiple comparisons test in C and by a paired t test in E. Scale bars = 10 μm.

We next analyzed the influence of Wnt signaling on myosin dynamics. Upon attachment to a bead and ABxx or P_2_ from Wnt mutants, myosin movements became asymmetric between B1 and B2 regions but not in U1 and U2 regions, leading to local flow asymmetries (Fig. 7C, D, G and Movie S8). However, attachment to a bead and a P_2_ cell yielded asymmetric myosin velocities for both U1-U2 and B1-B2 that generated global flow asymmetry (Fig. 7C, D, G, and Movie S8). Notably, myosin foci were always faster and had a longer lifetime in the cortex proximal to the P_2_ cell in the presence of Wnt (Figs. 7C, D and S6A and Movie S8), suggesting that Wnt instructively activates myosin activity.

To determine if Wnt activates myosin, we analyzed myosin II distribution and RLC phosphorylation. In both *in vivo* and *in vitro* experiments, localization of myosin II was diminished in the region proximal to P_2_, whereas that of p-RLC was more symmetric (Figs. 7E and S6B). Consequently, the level of p-RLC per myosin II—indicative of active myosin—was higher in the proximal cortex (Fig. 7E). Attachment to P_2_ cells from a Wnt mutant abolished myosin II asymmetry, causing myosin activity to become symmetric (Fig. 7E). Taken together, these results demonstrate that the Wnt signal is a cue that induces myosin activity asymmetry along an axis proximal-distal to Wnt-expressing cells. Unlike local myosin flow asymmetry induced by the physical contact, the Wnt-dependent global myosin flow asymmetry does not generate torque toward the contact site, probably because it moves cell surface of U1-U2 and B1-B2 regions in the same direction, resulting in orientation of EMS cell division toward P_2_ even in the presence of multiple other contact sites (Fig. 7G, orange arrows in the right panel).

## Discussion

In this study, we have discovered a cell division orientation mechanism that is independent of known microtubule-dynein pathways, using the well-established *C. elegans* model for cell division orientation but focusing on less studied cell types. We found that the 2-cell stage AB cell underwent oriented division independently of microtubule pulling forces. We identified physical contact with cells or bead acts as the upstream cue that orients AB cell division through downregulation of myosin activity. As a consequence, myosin equatorial flow became asymmetric in the region proximal to the contact site, while symmetric in the distal region, creating a local actomyosin flow asymmetry. By tracking cell surface movements, we showed that actomyosin flow is a force generator. Based on these data, we concluded that locally asymmetric cell surface movements generate a directional torque that orients cell division in parallel to the contact site. Notably, we found that a similar myosin-dependent mechanosensitive mechanism also operates in mouse 2-cell stage embryos. Furthermore, in later stage *C. elegans* embryos, additional cues such as contact asymmetry or Wnt signaling generate polarized actomyosin flows to specify a left-right oriented ABa cell division and an anterior-posterior oriented EMS cell division, respectively. Overall, our study demonstrates that cortical actomyosin flow is a cell-nonautonomously tunable force generation mechanism that orients cell division along different axes. In concert with microtubule pulling forces, tunable actomyosin flow may specify diverse division axis *in vivo*.

Our results provide the first examples of extrinsic control of actomyosin flow in dividing cells. The molecular mechanism of mechanosensitive actomyosin regulation in this system remains unclear. For example, we do not know how physical contact inhibits RLC phosphorylation. Extracellular control of actomyosin flow has also been reported in migrating cells, where a chemoattractant gradient acts through receptors to generate retrograde actomyosin flow toward the trailing end of cells (Devreotes and Horwitz, 2015). Rho GTPases such as Rac, Cdc42, and RhoA are actomyosin regulators critical for correct cell migration with spatio-temporally distinct activities within cells (Fritz and Pertz, 2016). As knock-down of RGA-3/RhoGAP resulted in the abnormal AB cell division, physical contact-induced myosin regulation may act through Rho GTPases in our system. However, we did not detect bead induced changes in RhoA and Cdc42 activities using GFP reporters that bind to their substrates, potentially due to their high cytoplasmic signals (data not shown; Kumfer et al., 2010; Tse et al., 2012). Development of cell surface specific reporters may allow more precise analysis. Alternatively, physical contact may directly inhibit myosin by exerting resistive forces; a previous single molecule study showed that purified myosin II molecules prematurely detached from F-actin and exhibited smaller working stroke when resistive forces were exerted (Capitanio et al., 2012). Notably, we showed that myosin lost the ability to respond to the contact site after knock-down of CUL-3 E3 ubiquitin ligase; thus future identification of CUL-3 substrates might shed light on the molecular details of mechanosensitive myosin regulation.

Since cells are usually surrounded by others from which they receive multiple cues, it is difficult to identify the cue that orients cell division in a multicellular system. Although reductionist approaches such as cell ablation or conditional protein knock-down play important roles in identifying cell-cell communications, reconstitution of multicellular physical and chemical environments enabled us to directly test the requirements for candidate cues. We have shown that cell-sized carboxylate-modified polystyrene beads are simple yet useful tools for examining the effects of physical contact on cell division. These chemically functionalized beads can also be modified with signaling molecules such as Wnt ligand (Habib et al., 2013). Although embryo micro-manipulations currently are demanding experiments, the future development of automated systems that spatially positions the cells with respect to properly designed beads would enable the systematic investigation of how different mechanical and chemical cues affect cell division in reconstituted three dimensional multicellular environments.

In this study, we uncovered the mechanism of 4-cell stage embryo architecture formation that is conserved between *C. elegans* and mouse, and potentially in human. These 4-cell stage architectures are significant for the further development of these animals: the diamond shape in *C. elegans* establishes the D-V axis, while the tetrahedron shape in mouse and human embryos promotes more successful *in vitro* fertilization outcomes than do other patterns (Cauffman et al., 2014; Ebner et al., 2012; Graham and Deussen, 1978; Suzuki et al., 1995) (Fig. 5C). Moreover, the tetrahedron and diamond shapes in mouse embryos are associated with distinct pluripotency factor activities and gene expression profiles among blastomeres (Goolam et al., 2016; Torres-Padilla et al., 2007; White et al., 2016). We also have shown that a mechanosensitive myosin pathway operates in both *C. elegans* and mouse embryos to orient cell division. In addition, this mechanism coordinately works with extracellular material such as eggshell or zona-pellucida to establish 4-cell stage architecture. Interestingly, zona-pellucida free 2-cell stage human embryos also appear to undergo cell division in parallel to cell contact (Bodri et al., 2015). Therefore, this newly identified oriented division mechanism may also function in human embryos. While extrinsic control is not known, actomyosin regulation at the 4-cell stage is associated with developmental delay or arrest in human embryos (Wong et al., 2010) and left-right body asymmetry in *C. elegans*, snail, and frog embryos (Natanathan et al., 2014; Davison et al., 2016). Thus, precise actomyosin regulation at the 4-cell stage may play central roles in early embryogenesis among Bilateria, and further studies of extrinsic actomyosin regulation would potentially contribute to our understanding of human multicellular assembly processes.

In a broader perspective, our results suggest that different patterns of cell contact between cells of distinct sizes and fates arising during embryogenesis function as unique mechanical and chemical cues that control actomyosin flow and cell division to produce diverse axis orientations. Indeed, *C. elegans* is remarkable in having an invariant lineage of division axis orientation (Sulston et al., 1983), and reproducible patterns of cell contact provide a potentially simple explanation for how this invariance is achieved. By examining the roles of each cell-cell contact on the orientation of unexplored cell divisions, future studies may uncover new types of cell division regulators, and reveal how sequences of contact pattern-dependent oriented divisions contribute to the generation of multicellular architecture throughout development.

## Author contributions

Conceptualization, K. S.; Methodology, K. S.; Formal Analysis, K. S.; Investigation, K. S.; Writing – Original Draft, K. S. and B. B.; Writing – Review & Editing, K. S. and B. B.; Visualization, K. S.; Funding Acquisition, K. S. and B. B.; Supervision, B. B.

## Acknowledgments

We thank Ute Hostick and University of Oregon transgenic mouse facility for providing mouse embryos, Paul Mains, Tony Hyman and the *Caenorhabditis* Genetics Center (funded by the NIH Office of Research Infrastructure Programs; P40 OD010440) for *C. elegans* strains, Akatsuki Kimura for helpful discussion, Kryn Stankunas and Chris Doe for sharing lab equipment, and Chris Doe for critical reading of the manuscript. This work was supported by the NIH grant R01GM049869 to B.B. and Human Frontier Science Program (LT000345/2012-L) to K.S.

---

**Figure S1.**
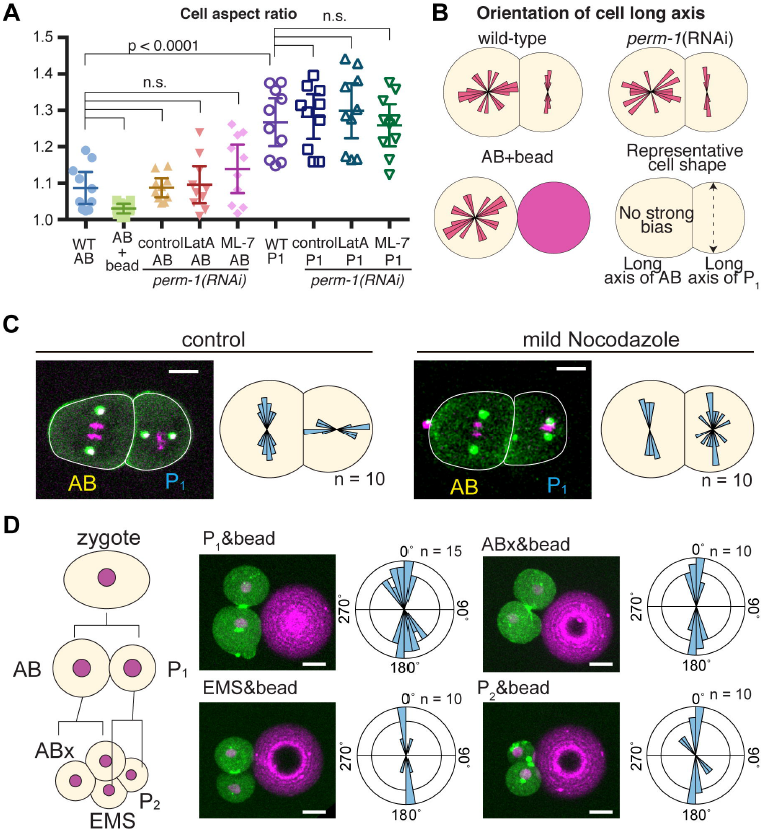
Related to Figure 1. Oriented AB division is independent of known microtubule-dynein pathways and other embryonic cells also respond to bead contact. (A) Cell aspect ratio of indicated cells. Except for isolated AB cell with bead, cells within eggshell were quantified. P-values were calculated by one-way ANOVA with Holm-Sidak’s multiple comparisons test. (B) Orientation of cell long axes. P_1_ cell has strong bias in transverse axis while AB is not biased. (C) AB and P_1_ division axes after treatment with 12. 5 ng/ml nocodazole. Centrosomes (green/white), histones (magenta), and cell outlines (white lines) are shown. Distributions of AB division axes at anaphase and those of P_1_ at metaphase are shown schematically. (D) Bead contact-induced oriented division in other cell types. The distribution of division axes after cytokinesis is shown as angular plots. Scale bars = 10 μm.

**Figure S2.**
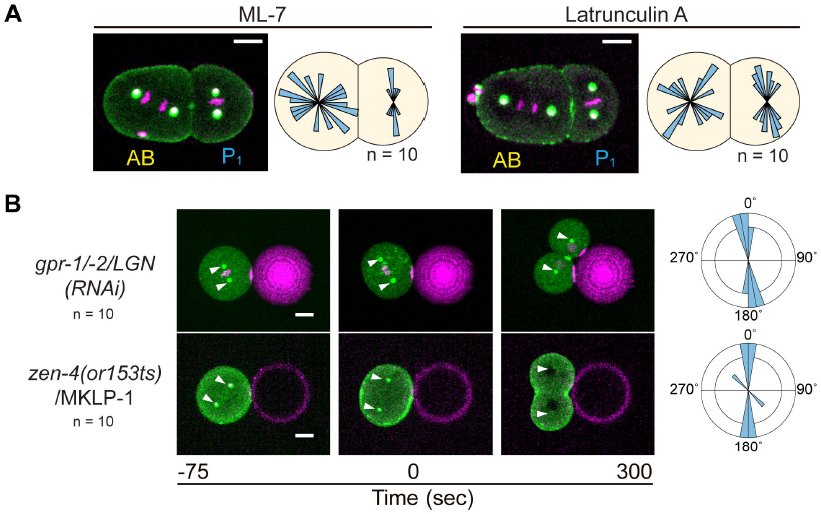
Related to Figure 2. Oriented AB division is actomyosin-dependent. (A) Randomized AB division axes after treatment with ML-7 or latrunculin A. The distribution of AB and P_1_ division axes is shown schematically. (B) Normal bead contact-dependent oriented AB division upon LGN knockdown or in mutants of cytokinesis. Arrowheads indicate centrosome positions. Angular plots indicate AB division axes after cell cycle exit (as evaluated by histone decondensation). Myosin (green), centrosomes (white/green), bead (magenta), and histones (magenta) are shown.

**Figure S3.**
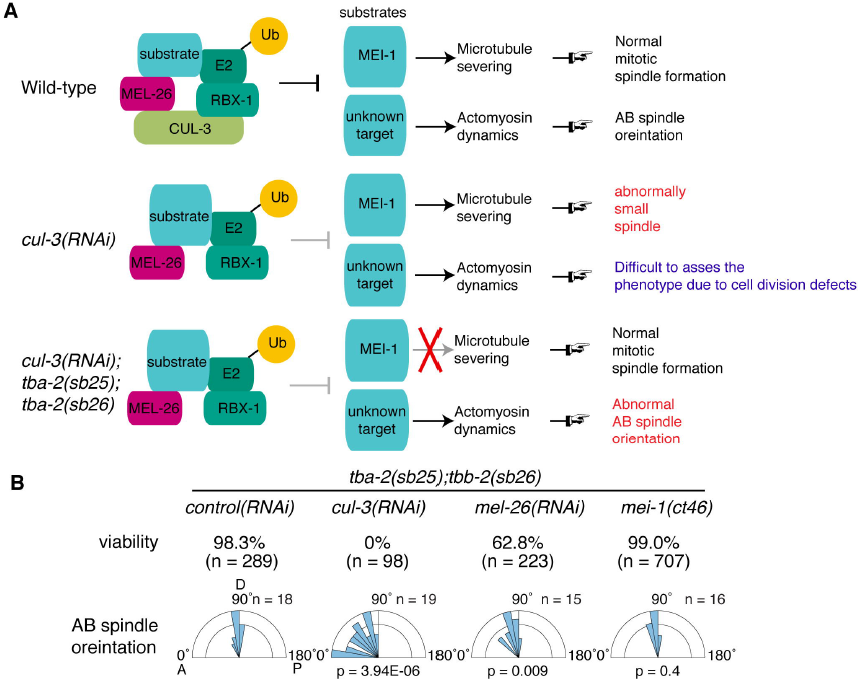
Related to Figure 2. CUL-3-MEL-26 E3 ubiquitin ligase complex regulates the AB division axis independent of MEI-1 function. (A) Schematic illustrations of genetic backgrounds used in this study. CUL-3/Cullin forms an E3 ubiquitin ligase complex with BTB protein MEL-26, RBX-1 and E2 enzyme to ubiquitylate substrate proteins. The CUL-3-MEL-26 complex regulates the degradation of the meiosis-specific microtubule-severing protein MEI-1 during the oocyte-to-embryo transition to allow formation of a normal mitotic spindle. Upon *cul-3* or *mel-26* knock-down or in degradation-defective *mei-1(ct46)* mutants, ectopic MEI-1 protein activity during mitosis causes the formation of abnormally small spindles with short microtubules (middle panel). However, when *tba-2(sb25)* and *tbb-2(sb26)* mutations are introduced (α-tubulin and β-tubulin alleles, respectively), microtubules become resistant to ectopic MEI-1 protein activity (Lu and Mains, 2005) and normal mitotic spindle formation is restored (bottom panel). In *cul-3*(RNAi) embryos, AB spindle orientation is defective in the *tba-2(sb25);tbb-2(sb26)* background, suggesting that this phenotype results from a failure in degradation of an unknown target. (B) Viability and AB spindle orientation after *cul-3* knockdown are independent of ectopic MEI-1 function. Eggs were laid by adult worms for around 6 h and hatched and dead embryos were counted to assess viability. Note that the degradation-defective mutant *mei-1(ct46)* showed 0% viability in a wild-type background (Lu and Mains, 2005), which was completely rescued to the wild-type level in the *tba-2(sb25);tbb-2(sb26)* background. Low viability and abnormal AB spindle orientation in *cul-3*(RNAi) and *mel-26*(RNAi) embryos suggest that CUL-3 and MEL-26 function as a complex to regulate the degradation of unknown targets other than MEI-1 protein. The weak phenotype in *mel-26*(RNAi) suggests that there are other BTB proteins involved in this process. A, P, and D in the graph indicate anterior, posterior, and dorsal sides, respectively. P values were calculated with the Watson-Williams test for equal means (n = 18, 19, 15, and 16 from left to right).

**Figure S4.**
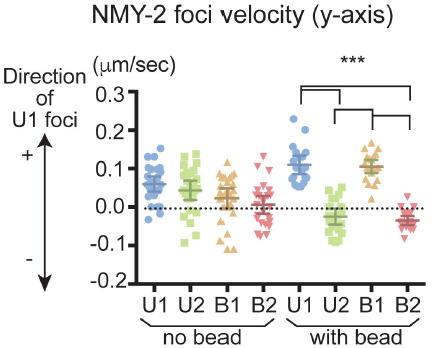
Related to Figure 3. Y axis myosin foci velocities after contact with a bead. Velocities were determined as in Figure. 3C. P values were calculated by one-way ANOVA with Holm-Sidak’s multiple comparisons test.

**Figure S5.**
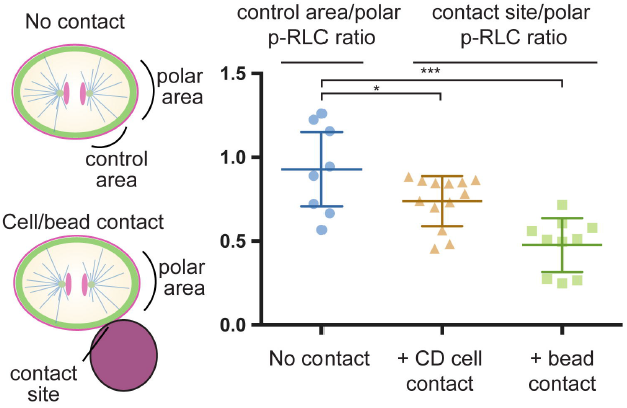
Related to Figure 5. Contact-dependent reduction of RLC phosphorylation. Left schematics indicate the cellular regions used for the quantification of phospho-RLC signals. In isolated cells, area between equator and pole are used as control area. Right graph shows the ratio of p-RLC signal between control and polar area in no contact and contact site and polar area in the presence of physical contact. P values were calculated by one-way ANOVA with Holm-Sidak’s multiple comparisons test.

**Figure S6.**
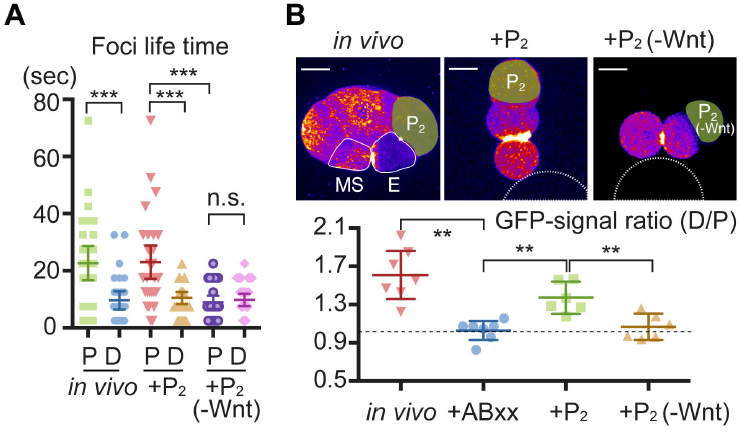
Related to Figure 7. Polarized myosin foci lifetime and intensity in the presence of Wnt signal. (A) Polarized myosin foci lifetime on the cortex of cell halves proximal and distal to the P_2_ cell. (B) Polarized myosin localization in the presence of Wnt. Heat maps of myosin GFP signal intensity are shown. White lines in the left panel indicate EMS cells and white dotted lines in the middle and right panels indicate beads. P values were calculated by one-way ANOVA with Holm-Sidak’s multiple comparisons test. *P < 0.05, **P < 0.01, ***P < 0.0001; n.s., not significant (P > 0.05). Scale bars = 10 μm.

**Movie S1.** Physical contact-dependent oriented AB cell division. Centrosomes, myosin (green), chromosomes (magenta), and beads (magenta) after cell-cell or cell-bead recombination (15 s per frame).

**Movie S2.** Physical contact-induced anisotropic myosin flow. Cortical myosin foci movements in a 3D reconstruction (left) or maximum projection with correction of cell rotation/movements (right) are shown for cells with or without bead attachment. Movie loops three times (5 s per frame in the movie on the right).

**Movie S3.** Equatorial actomyosin flow specifically becomes asymmetric proximal to contact. Cortical myosin foci movements were tracked in 3D and shown as ball. Left and right movies are from same sample showing distal and proximal side relative to the contact site, respectively.

**Movie S4.** Physical contact-induced anisotropic cell surface movements. Cortical myosin (green), beads (magenta, at the bottom of the image), and cell surface particles (magenta) after AB-bead contact. The movie loops three times (5 s per frame).

**Movie S5.** Physical contact-dependent oriented mouse AB/CD cell division. Differential interference contrast (DIC) movies of intact mouse embryos (left), zona-free embryos (left, marked with circles), and isolated AB/CD blastomeres attached to a bead (right) (5 min per frame).

**Movie S6.** Extracellular material-dependent cell rotation prevents perpendicular P_1_/CD cell division. DIC movies of an intact *C. elegans* embryo (upper left), eggshell-free *C. elegans* embryo (lower left), intact mouse embryo (upper right), and zona-free mouse embryo (lower right). In the presence of extracellular material, dividing *C. elegans* AB cells or in mouse CD cells rotates toward the cell contact.

**Movie S7.** Asymmetric-sized contacts induces polarized myosin flow. Surface myosin (green), chromosomes (magenta), and beads (magenta) are shown. Bottom panels show myosin movement after correction of cell movement/rotations. The movie loops three times (5 s per frame).

**Movie S8.** Wnt-induced polarized myosin flow. Surface myosin (green), chromosomes (magenta), and beads (magenta) are shown. Bottom panels show myosin movement after correction of cell movement/rotation. The movie loops three times (5 s per frame).

## STAR Methods

### Key Resource Table

Provided as a separate file

### Contact for Reagent and Resource Sharing

Lead contact information:

Kenji Sugioka, Institute of Molecular Biology, University of Oregon

Lead contact holds responsibility for responding to requests and providing reagents and information.

### Experimental Model and Subject Details

*Mus musculus* (mouse) and *Caenorhabditis elegans* strains were used in this study. To obtain mouse embryos, female C57BL/6J in 3-12 week old were superovulated by intraperitoneal injections of 5 international unit (IU) of Pregnant Mare Serum Gonadotropin (PMSG) followed by 5 IU of Human Chorionic Gonadotropin (hCG) 48 hours later. Each females was then placed with a C57BL/6J or B6.Cg-Tg(HIST1H2BB/EGFP)1Pa/J male (The Jackson Laboratory) overnight and all females were checked for a copulation plug the following morning. Two-cell stage embryos were collected by flushing oviducts with FHM medium and cultured in EmbryoMax Advanced KSOM medium (EMD Millipore) under 37°C, 5% CO_2_, and 5% O_2_. All *C. elegans* strains except for temperature sensitive mutants were cultured at 25°C as described (Brenner, 1974). *z*en-4(*or153*ts) temperature sensitive mutants were cultured at 15°C before experiments. The following transgenes were used: *ddIs299* (YFP::SPD-5, centrosome marker, a gift from Tony Hyman), *itIs37* (mCherry::histone H2B), *zuIs45* (NMY-2::GFP, non-muscle myosin II), *cp13*[*nmy-2*::GFP + LoxP] (non-muscle myosin II), *xsSi5*[GFP::*ani-1*(AH+PH); RhoA biosensor], and *ojIs40*[mGFP::*wsp-1*, CDC-42 biosensor]. Some strains carry the viable α-tubulin and β-tubulin gene mutations *tba-2(sb25)* and *tbb-2(sb26)* (Lu and Mains, 2005), respectively (gifts from Paul Mains), to suppress *cul-3(RNAi)* defects in early stage (see Figure S3).

## Method Details

### Suppression of early *cul-3* knock-down defects in worm embryos

CUL-3/Cullin and its adaptor binding partner MEL-26/BTB are E3 ubiquitin ligase components required for degradation of the meiosis-specific microtubule severing protein MEI-1/katanin, and *cul-3(RNAi)* results in abnormal mitotic spindles with short microtubules and associated early embryonic cell division defects, and embryonic lethality (Figure S3) (Kurz et al., 2002; Pintard et al., 2003). However, the suppressor mutations *tba-2(sb25)* and *tbb-2(sb26)* almost fully rescue the lethality and cell division defects of a *mei-1* dominant mutation *ct46*, which encodes a degradation-defective form of MEI-1 (Figure S3A and S3B) (Lu and Mains, 2005). *tba-2(sb25)* and *tbb-2(sb26)* confer resistance to MEI-1-dependent microtubule severing without affecting normal mitosis (Lu and Mains, 2005). We therefore carried out all experiments in *tba-2(sb25)*;*tbb-2(sb26)* mutant backgrounds to suppress early *cul-3(RNAi)* defects caused by ectopic MEI-1 function. Importantly, we confirmed that AB spindle orientation was normal in *mei-1(ct46); tba-2(sb25);tbb-2(sb26)* mutants, indicating that AB spindle orientation is regulated by the CUL-3-MEL-26 dependent degradation of unknown substrate(s) other than MEI-1 (Figure S3B). The weaker phenotype of *mel-26(RNAi);tba-2(sb25);tbb-2(sb26)* embryos suggests that other BTB proteins redundantly regulate the degradation of the unknown substrate(s).

### *C. elegans* RNAi

Feeding RNAi was performed for control, *cul-3, gpr-1/-2* and *perm-1* at 25°C. For control RNAi, a bacterial strain with empty vector (L4440) was used. For control and *gpr-1/-2* RNAi, L2 larvae were grown on the feeding RNAi plates and embryos were analyzed. For *cul-3* RNAi, L1 larvae were grown on the feeding RNAi plates and sterile F1 adults were crossed with RNAi males to obtain embryos for analysis. The sterile F1 adult worms have oocytes that appear normal but abnormally large sperm that probably failed to fully differentiate. For *perm-1* RNAi, L4 larvae were grown on the feeding RNAi plates for 10-14 hrs.

### Polystyrene beads preparation

10 mg carboxyl-modified polystyrene beads at 30 μm diameter (KISKER BIOTECH GmbH & Co.), 20 μm, 10 μm, and 0.35 μm diameters (Polysciences) were washed twice with 100 mM 2-(N-morpholino)ethanesulfonic acid (MES) buffer (pH 6.5) and incubated with the 1 mL MES buffer containing 10 mg 1-Ethyl-3-(3-dimethylaminopropyl) carbodiimide (EDAC) for 15 minutes at room temperature. The beads were washed twice with phosphate buffered saline (PBS) and incubated with the 0.5 mL PBS containing 0.05 μg Rhodamine Red-X succinimidyl ester (ThermoFisher Scientific) for 5 minutes. The beads were washed twice and stored with PBS at 4°C before use. Cells adhere to the beads either with or without poly-L-lysine coating, probably due to the interactions between carboxyl-modified bead and extracellular proteins or the electrostatic interactions between the negative charge of plasma membrane and positive charge of Rhodamine. Therefore, all experiments except Fig. 2A were done without poly-L-lysine coating.

### Blastomere isolation

For *C. elegans* embryos, embryonic cells were isolated as described before (Edgar, 1995; Park et al., 2005), with the following modifications. After cutting the adult worms in egg salt buffer, embryos were placed in the freshly made hypochlorite solution [75 % Clorox (Clorox) and 2.5 N KOH] for 50 seconds. After washing with Shelton’s growth medium twice (Shelton and Bowerman, 1996), embryos were put into the Shelton’s medium on the coverslip with metallic holds. Eggshell and permeabilization barrier were removed by repeated mouth pipetting with a hand-drawn glass microcapillary tubes (10 microliters, Kimble Glass Inc.) to obtain eggshell and permeabilization barrier-free embryos. Although the permeabilization barrier is more important for cell architectures than eggshell (Schierenberg and Junkersdorf, 1992), we call these embryos eggshell-free embryos for simplicity. For blastomere isolation, 2-cell stage eggshell-free embryos were further drawn to separate the two cells. ABx, ABxx, EMS and P_2_ cells were isolated by separating daughter cells after each cell division.

Two-cell stage mouse embryos were placed in M2 medium containing Tyrode’s acid (Sigma-Aldrich) briefly to remove zona-pellucida. They were used as zona-free embryos or further separated by repeated mouth pipetting with a hand-drawn glass microcapillary tubes to obtain individual blastomere.

### Cell-beads interaction assay

Isolated cells at least before prometaphase were attached to either one or two polystyrene beads before imaging. For 1cell-2 beads assay, 2 beads were combined initially and then attached to the isolated cell. For the surface particle tracking, cells were first coated with 0.35 μm beads and then attached to the 30 μm bead. All manipulations were done with the mouth pipette. For *zen-4* temperature sensitive mutants, isolated AB cells were prepared at 15 °C and incubated at 25 °C using a temperature controlled stage equipped with Leica DMi8 microscope (Leica) described below.

### Drug treatments

For the drug treatments of *C. elegans* embryos surrounded by the eggshell (intact embryo), we performed *perm-1* RNAi to permeabilize the eggshell as previously described (Carvalho et al., 2011). Early 2-cell stage embryos were transferred to the Shelton’s growth medium containing 2% DMSO (control), 20 μM Latrunculin A (Sigma-Aldrich), 200 μM ML-7, 12.5 ng/mL Nocodazole (Sigma-Aldrich) and 20 μg/ml Nocodazole.

For the drug treatments of *C. elegans* isolated blastomere, Shelton’s growth medium containing 200 μM ML-7 (Sigma-Aldrich) and 20 μg/ml Nocodazole (Sigma-Aldrich) were used. For drug treatments of mouse blastomere, EmbryoMax Advanced KSOM medium containing 50 μM ML-7 was used.

### Live-imaging

For worm cells with eggshell removed or permeabilized, metallic slides with a hole in the center were sandwiched between two coverslips and cells were mounted on one of the coverslip with Shelton’s medium to avoid compression of the cell and dehydration. For other *C. elegans* experiments, embryos were mounted on the 4% agar on the slide glass and sealed with a petroleum jelly (Vaselline) after placing coverslips. For mouse embryos, embryos in a drop of EmbryoMax Advanced KSOM medium covered with mineral oil on cell culture dish with glass bottom (FluoroDish, World Precision Instruments) were used for the imaging. For worm DIC movies, imaging was performed with generic CCD cameras mounted on either AxioPlan or AxioSkop (Zeiss). For worm fluorescence imaging, imaging was performed with a confocal unit CSU10 (Yokogawa electric) and an EMCCD camera Image EM (Hamamatsu photonics) mounted on an inverted microscope Leica DMI 4000 (Leica) or performed with a confocal unit CSU-W with Borealis (Andor) and an EMCCD camera iXon Ultra 897 (Andor) mounted on an inverted microscope Leica DMi8 (Leica). Both imaging systems were controlled by Metamorph (Molecular Devices). Samples were illuminated by diode-pumped lasers with 488 nm and 561nm wavelength. NMY-2::GFP, YFP::SPD-5 and mCherry::histone H2B signals were obtained every 15 sec with 2 μm Z spacing for most experiments. NMY-2::GFP, mCherry::histone H2B and 0.35 μm particle signals were obtained every 5 sec with 1.5 μm Z spacing for the tracking of cortical movements in cell-bead experiments. NMY-2::GFP, mCherry::histone H2B signals were obtained every 10 sec with 1.5 μm Z spacing for the tracking of cortical NMY-2 movements in intact embryos. For the live-imaging of mouse embryo, Nikon Eclipse Ti inverted fluorescence microscope (Nikon) equipped with Live Cell stage top incubation system (Pathology Devices) controlled by NIS Elements Advanced Research software was used. Embryos were maintained at 37°C, 5% CO_2_, and 5% O_2_ condition.

### Immunofluorescence

Mouse or *C. elegans* embryos were fixed with 4% Paraformaldehyde in PBS for 15 mins. Fixed embryos were washed three times with PBS containing 1% Tween-20. Embryos were then incubated with anti-phospho-myosin light chain (Ser19) antibody (1:50 dilution, #3671, Cell Signaling Technology) in PBS containing 1% Bovine Serum Albumin (BSA) and 0.1% Tween-20 at 4°C for overnight. After washing three times, samples were incubated with 1:500 anti-rabbit Rhodamine Red-X (Jackson ImmunoResearch) for 2hrs at room temperatures. Mouse embryos were imaged at 0.5 μm Z spacing while those of worms were at 0.25 μm.

### Image analysis

All images were analyzed with Fiji (Schindelin et al., 2012). For Fig. 1B and Fig. 2A, the orientations of mitotic spindles were calculated from the angles between the line that crosses the longitudinal axis (A-P) of the embryo and either AB or P_1_ spindle using DIC movies. As AB cell starts to rotate after late anaphase due to the eggshell, and also affects P_1_ spindle orientation, we measured the spindle orientations before the cell rotation. Therefore, AB spindle at anaphase and P_1_ spindle at metaphase were measured. D-V axes were measured from the angle between (i) the line that passes the centroids of ABp and EMS nuclei at 4-cell stage and (ii) the A-P axis, using DIC movies. For other experiments, spindle orientations relative to the plane of cell-bead or cell-cell contact at indicated cell cycle stages were measured using the 3D reconstructed fluorescence images. All angular data were analyzed with PAST software (Hammer et al., 2001). For the quantification of myosin foci movements in the cell, cell position and orientation were corrected by the image J plug-in StackReg, to eliminate the movements of myosin foci caused by the cell movements or rotation. Supplemental Movies were made with Adobe Premiere Elements 15 (Adobe) or Imaris (Bitplane).

### Quantification and Statistical Analysis

Error bars indicate the mean ± 95% confidence interval. Statistical methods were described in the Figure legends. Statistical analysis was performed using PAST software, Microsoft Excel (Microsoft) and Prism 6 (GraphPad Software). *P < 0.05, **P < 0.01, ***P < 0.0001; n.s., not significant (P > 0.05).

## References

Adelstein, R. S. and Conti, A. M. (1975). Phosphorylation of platelet myosin increases actin-activated myosin ATPase activity. Nature 256, 597-598.

Basto, R., Lau, J., Vinogradova, T., Gardiol, A., Woods, C. G., Khodjakov, A., Raff, J. W. (2006). Flies without centrioles. Cell 125, 1375-1386.

Bazzi, H., Anderson, K. V. (2014). Acentriolar mitosis activates a p53-dependent apoptosis pathway in the mouse embryo. Proc. Natl. Acad. Sci. U. S. A. 111, E1491-E1500.

Bodri, D., Kato, R., Kondo, M., Hosomi, N., Katsumata, Y., Kawachiya, S., Matsumoto, T. (2015). Time-lapse monitoring of zona pellucida-free embryos obtained through *in vitro* fertilization: a retrospective case series. Fertil. Steril. 103, e35.

Bosveld, F., Markova, O., Martin, C., Wang, Z., Pierre, A., Gaugue, I., Ainslie, A., Christophorou, N., Lubensky, D. K., Minc, N., Bellïche, Y. (2016). Epithelial tricellular junctions act as interphase cell shape sensors to orient mitosis. Nature 530, 495-498.

Bray, D. and White, J. G. (1988). Cortical flow in animal cells. Science 239, 883-888.

Brenner, S. (1974). The genetics of *Caenorhabditis elegans*. Genetics 77, 71-94.

Campinho, P., Behrndt, M., Ranft, J., Risler, T., Minc, N., Heisenberg, C. P. (2013). Tension-oriented cell divisions limit anisotropic tissue tension in epithelial spreading during zebrafish epiboly. Nat. Cell Biol. 15, 1405-1411.

Capitanio, M., Canepari, M., Maffei, M., Beneventi, D., Monico, C., Vanzi, F., Bottinelli, R., Pavone, F. S. (2012). Ultrafast force-clamp spectroscopy of single molecules reveals load dependence of myosin working stroke. Nat. Methods 9, 1013-1019.

Carvalho, A., Olson, S. K., Gutierrez, E., Zhang, K., Noble, L. B., Zanin, E., Desai, A., Groisman, A., Oegema. K. (2011). Acute drug treatment in the early *C. elegans* embryo. PloS One 6, e24656.

Case, L. B. and Waterman, C. M. (2015). Integration of actin dynamics and cell adhesion by a three-dimensional, mechanosensitive molecular clutch. Nat. Cell Biol. 17, 955-963.

Cauffman, G., Verheyen, G., Tournaye, H., Van de Velde, H. (2014). Developmental capacity and pregnancy rate of tetrahedral-versus non-tetrahedral-shaped 4-cell stage human embryos. J. Assist. Reprod. Genet. 31, 427-34.

Craig, R., Smith, R., Kendrick-Jones, J. (1983). Light-chain phosphorylation controls the conformation of vertebrate non-muscle and smooth muscle myosin molecules. Nature 302, 436-439.

Devreotes, P. and Horwitz, A.R. (2015). Signaling networks that regulate cell migration. Cold Spring Harb. Perspect. Biol. 7, a005959.

Davison, A., McDowell, G. S., Holden, J. M., Johnson, H. F., Koutsovoulos, G. D., Liu, M. M., Hulpiau, P., Van Roy, F., Wade, C. M., Banerjee, R., et al. (2016). Formin Is Associated with Left-Right Asymmetry in the Pond Snail and the Frog. Curr. Biol. 26, 654-660.

di Pietro, F., Echard, A., Morin, X. (2016). Regulation of mitotic spindle orientation: an integrated view. EMBO Rep. 17, 1106-30.

Ebner, T., Maurer, M., Shebl, O., Moser, M., Mayer, R. B., Duba, H. C., Tews, G. (2012). Planar embryos have poor prognosis in terms of blastocyst formation and implantation. Reprod. Biomed. Online 25, 267-72.

Edgar, L. G. (1995). Blastomere culture and analysis. Method Cell Biol 48, 303-321.

Egger, B., Boone, J. Q., Stevens, N. R., Brand, A. H., Doe, C. Q. (2007). Regulation of spindle orientation and neural stem cell fate in the *Drosophila* optic lobe. Neural Dev. 2, 1.

Fink, J., Carpi, N., Betard, A., Chebah, M., Azioune, A., Bornens, M., Sykes, C., Fetler, L., Cuvelier, D., Piel, M. (2011). External forcs control mitotic spindle positioning. Nat. Cell Biol. 13, 771-778.

Fritz, R. D., Pertz, O. (2016). The dynamics of spatio-temporal Rho GTPase signaling: formation of signaling patterns. F1000Research 5, 749.

Goolam, M., Scialdone, A., Graham, S. J., Macaulay, I. C., Jedrusik, A., Hupalowska, A., Voet, T., Marioni, J. C., Zernicka-Goetz, M. (2016). Heterogeneity in Oct4 and Sox2 Targets Biases Cell Fate in 4-Cell Mouse Embryos. Cell 165, 61-74.

Goldstein, B., Takeshita, H., Mizumoto, K., Sawa, H. (2006). Wnt signals can function as positional cues in establishing cell polarity. Dev. Cell 10, 391-396.

Graham, C. F. and Deussen, Z. A. (1978). Features of cell lineage in preimplantation mouse development. J. Embryol. Exp. Morph. 48, 53-72.

Habib, S. J., Chen, B. C., Tsai, F. C., Anastassiadis, K., Meyer, T., Betzig, E., Nusse, R. (2013). A localized Wnt signal orients asymmetric stem cell division *in vitro*. Science 339, 1445-1448.

Hammer, Ø., Harper, D. A. T., Ryan, P. D. (2001). PAST: Paleontological statistics software package for education and data analysis. Palaeontol. Electron. 4, 9pp, http://palaeo- electronica.org/2001_1/past/issue1_01.htm.

Hao, Y., Du, Q., Chen, X., Zheng, Z., Balsbaugh, J. L., Maitra, S., Shabanowitz, J., Hunt, D. F., Macara, I. G. (2010). Par3 controls epithelial spindle orientation by aPKC-mediated phosphorylation of apical Pins. Curr. Biol. 20, 1809-1818.

Kelly, S.J., Mulnard, J.G., and Graham, C.F. (1978). Cell division and cell allocation in early mouse development. J. Embryol. Exp. Morphol. 48, 37-51.

Knoblich, J. (2010). Asymmetric cell division: recent developments and their implications for tumour biology. Nat Rev Mol Cell Biol 11, 849-860.

Kumfer, K. T., Cook, S. J., Squirrell, J. M., Eliceiri, K. W., Peel, N., O’Connell, K. F., White, J. G. (2010). CGEF-1 and CHIN-1 regulate CDC-42 activity during asymmetric division in the *Caenorhabditis elegans* embryo. Mol. Biol. Cell 21, 266-277.

Kurz, T., Pintard, L., Willis, J. H., Hamill, D. R., Gönczy, P., Peter, M., Bowerman, B. (2002). Cytoskeletal regulation by the Nedd8 ubiquitin-like protein modification pathway. Science 295, 1294-1298.

Kwon, M., Bagonis, M., Danuser, G., Pellman, D. (2015). Direct Microtubule-Binding by Myosin-10 Orients Centrosomes toward Retraction Fibers and Subcortical Actin Clouds. Dev. Cell 34, 323-337.

Levayer, R. and Lecuit, T. (2012). Biomechanical regulation of contractility: Spatial control and dynamics. Trends Cell Biol. 22, 61-81.

Louvet-Vallee, S.,Vinot, S., Maro, B. (2005). Mitotic spindles and cleavage planes are oriented randomly in the two-cell mouse embryo. Curr. Biol. 15, 464-9.

Lu, C. and Mains, P. E. (2005). Mutations of a redundant α-tubulin gene affect *Caenorhabditis elegans* early embryonic cleavage via MEI-1/katanin-dependent and - independent pathways. Genetics 170, 115-126.

Mayer, M., Depken, M., Bois, J. S., Jülicher, F., Grill, S. W. (2010). Anisotropies in cortical tension reveal the physical basis of polarizing cortical flows. Nature 467, 617-621.

Minc, N., Burgess, D., Chang, F. (2011). Influence of cell geometry on division-plane positioning. Cell 144, 414-426.

Murthy, K. and Wadsworth, P. (2005). Myosin-II-dependent localization and dynamics of F-actin during cytokinesis. Curr. Biol. 15, 724-731.

Naganathan, S. R., Furthauer, S., Nishikawa, M., Julicher, F., Grill, S. W. (2014). Active torque generation by the actomyosin cell cortex drives left-right symmetry breaking. eLife 3, e04165.

Nestor-Bergmann, A., Goddard, G., Woolner, S. (2014). Force and the spindle: mechanical cues in mitotic spindle orientation. Semin. Cell Dev. Biol. 34, 133-139.

Noatynska, A., Gotta, M., Meraldi, P. (2012). Mitotic spindle (DIS) orientation and DISease: Cause or consequence? J. Cell Biol. 199, 1025-1035.

Park, F. D. and Priess, J. R. (2003). Establishment of POP-1 asymmetry in early *C. elegans* embryos. Development 130, 3547-3556.

Pease, J. C. and Tirnauer, J. S. (2011). Mitotic spindle misorientation in cancer—out of alignment and into the fire. J. Cell Sci. 124, 1007-1016.

Pintard, L., Willis, J. H., Willems, A., Johnson, J. L., Srayko, N., Kurz, T., Glaser, S., Mains, P. E., Tyers, M., Bowerman, B., Peter, M. (2003). The BTB protein MEL-26 is a substrate-specific adaptor of the CUL-3 ubiquitin-ligase. Nature 425, 311-316.

Priess, J. R. and Thomson, J. N. (1987). Cellular interactions in early *C. elegans* embryos. Cell 48, 241-250.

Priess, J. R. (2005). Notch signaling in the *C. elegans* embryo. WormBook, ed. The C. elegans Research Community, WormBook, doi:10.1895/wormbook.1.4.1

Reymann, A. C., Staniscia, F., Erzberger, A., Salbreux, G., Grill, S. W. (2016). Cortical flow aligns actin filaments to form a furrow. eLife 5, e17807.

Rocheleau, C. E., Downs, W. D., Lin, R., Wittmann, C., Bei, Y., Cha, Y. H., Ali, M., Priess, J. R., Mello, C. C. (1997). Wnt signaling and an APC-related gene specify endoderm in early *C. elegans* embryos. Cell 90, 707-716.

Rose, L., Gönczy, P. (2014) Polarity establishment, asymmetric division and segregation of fate determinants in early *C. elegans* embryos. WormBook, ed. The C. elegans Research Community, WormBook, doi:10.1895/wormbook.1.30.2

Roh-Johnson, M., Shemer, G., Higgins, C. D., McClellan, J. H., Werts, A. D., Tulu, U. S., Gao, L., Betzig, E., Kiehart, D. P., Goldstein, B. (2012). Triggering a cell shape change by exploiting preexisting actomyosin contractions. Science 335, 1232-1235.

Saitoh, M., Ishikawa, T., Matsushima, S., Naka, M., Hidaka, H. (1987). Selective inhibition of catalytic activity of smooth muscle myosin light chain kinase. J. Biol. Chem. 262, 7796– 7801.

Sawa, H., Korswagen H. C. (2013). Wnt signaling in *C. elegans*. WormBook, ed. The C. elegans Research Community, Wormbook, doi:10.1895/wormbook.1.7.2

Schierenberg, E. and Junkersdorf, B. (1992). The role of eggshell and underlying vitelline membrane for normal pattern formation in the early *C. elegans* embryo. Rouxs. Arch. Dev. Biol. 202, 10-16.

Schindelin, J., Arganda-Carreras, I., Frise, E., Kaynig, V., Longair, M., Pietzsch, T., Preibisch, S., Rueden, C., Saalfeld, S., Schmid, B., Tinevez, J. Y., White, D. J., Hartenstein, V., Eliceiri, K., Tomancak, P., Cardona, A. (2012). Fiji: an open-source platform for biological-image analysis. Nat. Methods 9, 676-682.

Schlesinger, A., Shelton, C., Maloof, J., Meneghini, M., Bowerman, B. (1999). Wnt pathway components orient a mitotic spindle in the early *Caenorhabditis elegans* embryo without requiring gene transcription in the responding cell. Genes Dev. 13, 2028-2038.

Scholey, J. M., Taylor, K. A., Kendrick-Jones, J. (1980). Regulation of non-muscle myosin assembly by calmodulin-dependent light chain kinase. Nature 287, 233-235.

Schonegg, S., Constantinescu, A. T., Hoege, C., Hyman, A. A. (2007). The Rho GTPase-activating proteins RGA-3 and RGA-4 are required to set the initial size of PAR domains in *Caenorhabditis elegans* one-cell embryos. Proc. Natl. Acad. Sci. 104, 14976-14981.

Seldin, L., Poulson, N. D., Foote, H. P., Lechler, T. (2013). NuMA localization, stability, and function in spindle orientation involve 4.1 and Cdk1 interactions. Mol. Biol. Cell 24, 3651– 3662.

Shelton, C. A. and Bowerman, B. (1996). Time-dependent responses to *glp-1*-mediated inductions in early *C. elegans* embryos. Development 122,2043-2050.

Siller, K. H. and Doe, C. Q. (2009). Spindle orientation during asymmetric cell division. Nat. Cell. Biol. 11 365-374.

Singh, D. and Pohl, C. (2014). Coupling of rotational cortical flow, asymmetric midbody positioning, and spindle rotation mediates dorsoventral axis formation in *C. elegans*. Dev. Cell 28, 253-267.

Sonnichsen, B., et al. (2005). Full-genome RNAi profiling of early embryogenesis in *Caenorhabditis elegans*. Nature 434, 462-9.

Srinivasan, D. G., Fisk, R. M., Xu, H., van den Heuvel, S. (2003). A complex of LIN-5 and GPR proteins regulates G protein signaling and spindle function in C. elegans. Genes Dev. 17, 1225-1239.

Sulston, J. E. and Horvitz, H. R. (1977). Post-embryonic cell lineages of the nematode, Caenorhabditis elegans. Dev. Biol. 56, 110-156.

Sulston, J. E., Schierenberg, E., White, J. G., and Thomson, J. N. (1983). The embryonic cell lineage of the nematode *Caenorhabditis elegans*. Dev. Biol. 100, 64-119.

Suzuki, H., Togashi, M., Adachi, J., Toyoda, Y. (1995). Developmental ability of zona-free mouse embryos is influenced by cell association at the 4-cell stage. Biol. Reprod. 53, 78-83.

Thorpe, C. J., Schlesinger, A., Carter, J. C., Bowerman, B. (1997). Wnt signaling polarizes an early *C. elegans* blastomere to distinguish endoderm from mesoderm. Cell 90, 695-705.

Torres-Padilla, M. E., Parfitt, D. E., Kouzarides, T., Zernicka-Goetz, M. (2007). Histone arginine methylation regulates pluripotency in the early mouse embryo. Nature 445, 214-8.

Tse, Y. C., Werner, M., Longhini, K. M., Labbe, J. C., Goldstein, B., Glotzer, M. (2012). RhoA activation during polarization and cytokinesis of the early *Caenorhabditis elegans* embryo is differentially dependent on NOP-1 and CYK-4. Mol. Biol. Cell 23, 4020-4031.

Uehara, R., Goshima, G., Mabuchi, I., Vale, R. D., Spudich, J. A., Griffis, E. R. (2010). Determinants of myosin II cortical localization during cytokinesis. Curr. Biol. 20, 1080– 1085.

Vicente-Manzanares, M., Ma, X., Adelstein, R.S., Horwitz, A.R. (2009). Non-muscle myosin II takes centre stage in cell adhesion and migration. Nat. Rev. Mol. Cell Biol. 10, 778-790.

White, M. D., Angiolini, J. F., Alvarez, Y. D., Kaur, G., Zhao, Z. W., Mocskos, E., Bruno, L., Bissiere, S., Levi, V., Plachta, N. (2016). Long-Lived Binding of Sox2 to DNA Predicts Cell Fate in the Four-Cell Mouse Embryo. Cell 165, 75-87.

Williams, S.E., Fuchs, E. (2013) Oriented divisions, fate decisions. Curr Opin Cell Biol 25, 749-758.

Williams, S. E., Ratliff, L. A., Postiglione, M. P., Knoblich, J. A., Fuchs, E. (2014). Par3-mInsc and Gαi3 cooperate to promote oriented epidermal cell divisions through LGN. Nat. Cell Biol. 16, 758-769.

Wilson, E. B. (1925). The Cell in Development and Heredity. (New York: The MacMillan Company).

Wong, C. C., Loewke, K. E., Bossert, N. L., Behr, B., De Jonge, C. J., Baer, T. M., Pera, R. A. R. (2010). Non-invasive imaging of human embryos before embryonic genome activation predicts development to the blastocyst stage. Nat. Biotech. 28, 1115-1121.

Wu, M., Smith, C. L., Hall, J. A., Lee, I., Luby-Phelps, K., Tallquist, M. D. (2010). Epicardial spindle orientation controls cell entry into the myocardium. Dev. Cell 19, 114-125.

Yamamoto, K., Kimura, A. (2017). An anisropic attraction model for the diversity and robustness of cell arrangement in nematodes. bioRxiv 132654. doi: https://doi.org/10.1101/132654

Yumura, S., Ueda, M., Sako, Y., Kitanishi-Yumura, T., Yanagida, T. (2008). Multiple mechanisms for accumulation of myosin II filaments at the equator during cytokinesis. Traffic 9, 2089-2099.

Zhou, M. and Wang, Y. L. (2008). Distinct pathways for the early recruitment of myosin II and actin to the cytokinetic furrow. Mol. Biol. Cell 19, 318-326.

Zonies, S., Motegi, F., Hao, Y., Seydoux, G. (2010). Symmetry breaking and polarization of the *C. elegans* zygote by the polarity protein PAR-2. Development 137, 1669-1677.

